# Microbial activation converts neutrophils into anti-tumor effectors

**DOI:** 10.1101/2020.08.21.259051

**Authors:** Andrew O. Yam, Jacqueline Bailey, Francis Lin, Arnolda Jakovija, Claudio Counoupas, James A. Triccas, Matthias Gunzer, Tobias Bald, Shane T. Grey, Tatyana Chtanova

## Abstract

Neutrophils infiltrate most solid tumors and their presence is usually correlated with suppression of anti-tumor responses, metastasis and poor prognosis. Here we used microbial bioparticles administered into the tumor microenvironment to transform neutrophils into anti-tumor effectors. Microbially activated neutrophils acquired an effector phenotype associated with pathogen killing and lost vascular endothelial growth factor expression associated with tumor growth and metastasis. They became the dominant immune cell infiltrating the tumor and inhibited tumor growth. Using intravital two-photon microscopy microbially activated neutrophils could be seen forming close contacts with tumor cells resulting in tumor tissue remodeling and tumor cell death. Thus, microbial bioparticle treatment can endow neutrophils with anti-tumor properties, suggesting that neutrophil plasticity in cancer could be exploited for tumor killing. These data highlight a pathway for the rational development of neutrophil-based cancer therapy.

## INTRODUCTION

Immunotherapies, especially T cell-centered approaches such as checkpoint inhibitors, have revolutionized the treatment of cancer (1). However, therapies that enhance antitumor responses by activating cytotoxic T cells have only been effective in a subset of cancer patients (2). Accordingly, there is an unmet clinical need for therapeutic approaches targeting other immune subsets. Beside strategies to harness the adaptive immune system, recent research highlighted immunotherapeutic potential of innate immune cells, including macrophages, dendritic and natural killer cells (3). Despite these, neutrophils, the most abundant immune subset with a vital role in microbial defense, remain largely overlooked as a target of cancer immunotherapy.

Neutrophils present the first line of protection against infection and are critical to our daily survival (4). They possess an extensive array of intracellular machinery that destroys microorganisms through phagocytosis, immune cell recruitment, release of antimicrobial molecules and tissue remodeling (5). This powerful killing potential of these cells is an outstanding example of their ability to fight disease (6).

In cancer neutrophils infiltrate most solid tumors (7), where they can promote tumor growth by stimulating tumor angiogenesis and contributing to the suppressive microenvironment within tumors *via* production of cytokines and chemokines that limit productive immune responses (8). Neutrophil capacity to suppress anti-tumor responses led them to be termed Myeloid-Derived Suppressor Cells (MDSCs) (7). Tumorinfiltrating neutrophils can also aid metastasis (9–11) and impair the effectiveness of antitumor immune therapies (12–15). Neutrophil presence in tumors is, therefore, frequently associated with a poor prognosis (7).

However, human and animal studies have also identified beneficial roles for neutrophils in cancer, such as direct cytotoxicity towards tumor cells and attenuation of metastasis formation (7). Like other cells of the immune system, neutrophils appear to rely on external cues to determine their activation and phenotypic state. Thus, anti-tumor and tumor-promoting neutrophil populations have been described (16).

Neutrophil function can be modulated depending on the factors present in the tumor microenvironment (TME) to either promote or oppose cancer progression, suggesting that manipulating neutrophil plasticity in cancer can be of therapeutic benefit. However, research on these terminally differentiated cells has been hampered by their experimentally intractable nature: neutrophils do not proliferate, are difficult to maintain in culture and are easily activated *ex vivo*. Furthermore, little is known about neutrophil persistence within tumors and how their function changes in response to signals within the TME. Delineation of neutrophil behavior in different settings using *in vivo* approaches is vital to our understanding of the precise role of these cells in tumor growth and metastasis as well as determining their potential as targets for cancer immunotherapy. Here, we demonstrate that bacterial stimulation can be used to recruit tumor neutrophils with a specific microbe-activated (MA) phenotype characterized by increased activation, enhanced motility and cytotoxic capacity. Using techniques such as intravital microscopy and photoconversion, we define neutrophil phenotype and function in cancer and show how these can be manipulated to endow neutrophils with anti-tumor effector capabilities and capacity to destroy tumor cells.

## RESULTS

### Microbial bioparticle treatment recruits and activates tumor neutrophils

We hypothesized that by introducing bacterial particles into tumors, it is possible to generate MA neutrophils with a distinct phenotype and anti-tumor functions. To test this, we injected *Staphylococcus aureus* (*S. aureus*) bioparticles directly into Lewis Lung Carcinoma (LLC) tumors established in the ear pinnae of C57BL/6 mice. We found that 24 hours after microbial bioparticle treatment there was a striking increase in immune cell infiltration within tumors (**Fig. 1A**).

**Figure 1.**
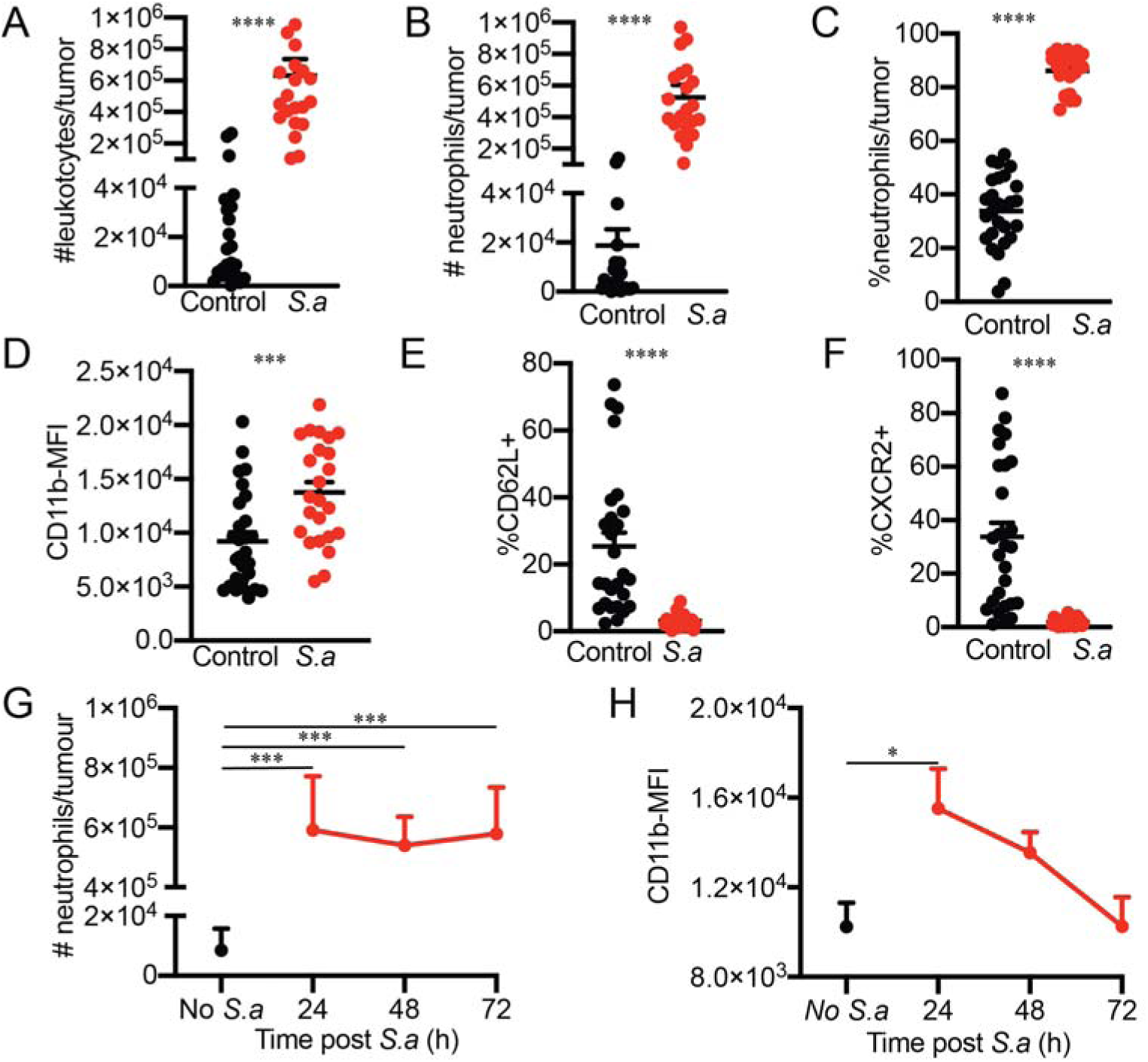
Microbe-driven recruitment and activation of intratumoral neutrophils. *S. aureus* bioparticle injection into tumors leads to an influx of leukocytes (A) and particularly neutrophils (B) which made up the majority of leukoctyes (C). Neutrophil activation was demonstrated by upregulation of neutrophil activation marker CD11b (D) and downregulation of CD62L (E) and CXCR2 (F) assessed by flow cytometry 24 hours post *S. aureus* injection. Neutrophil number (G) and CD1 lb expression (H) over time after an injection of 5. *aureus* bioparticles. Mean + SEM from five (A-F) and two (G-H) independent experiments. Each circle (A-D) represents a tumor. Data analyzed using non-parametric Mann-Whitney test (A-B, E), t-test (C-D) and Kruskal-Wallis test (G-H). * P ≤ 0.05 *** p ≤ 0.001 **** P ≤ 0.0001.

Neutrophils (identified as Ly6G^+^CD11b^+^ cells) were responsible for most of this increase as their number in tumors grew 28-fold (**Fig. 1B**) and comprised >85% of the tumor immune infiltrate following microbial bioparticle treatment (**Fig. 1C**). We also observed increased recruitment of macrophages and dendritic cells (**Supplementary Fig. 1A**), as well as recruitment and activation of T cells (**Supplementary Fig. 1B and C**). Our analysis indicated that CD4^−^CD8^−^ T cells were responsible for most of the increase in T cell number and activation. These results demonstrate that microbial bioparticle treatment dramatically re-shapes the tumor immune landscape driving strong recruitment of immune cells, and especially neutrophils, into tumors.

We next investigated the phenotype of tumor-infiltrating neutrophils and found that microbial bioparticle treatment led to an increase in the expression of neutrophil activation marker CD11b (**Fig. 1D**) and concomitant downregulation of CD62L (**Fig. 1E**) and CXCR2 (**Fig. 1F**), a profile consistent with neutrophil activation. This provides evidence that microbial treatment alters the phenotype of intratumoral neutrophils.

After the initial rapid influx of activated neutrophils, neutrophil numbers in tumors remained stable over the course of 72 hours (**Fig. 1G**). However, neutrophil activation slowly decreased and by 48 hours after treatment was similar to control tumors. This indicates that over time signals within the TME downmodulate neutrophil activation induced by our microbial bioparticle treatment and suggests that restimulation is required to maintain neutrophil activation (**Fig. 1H**).

### Microbial bioparticle treatment inhibits tumor growth

Microbial bioparticle treatment reshaped the tumor immune landscape leading to a substantial influx of MA neutrophils suggesting that this may present a promising approach to treat tumors. To test this hypothesis, we compared tumor growth in C57BL/6 mice with LLC tumors that were treated with microbial bioparticles to tumor-bearing mice that did not receive microbial treatment (**Fig. 2A**). We observed a striking decrease in tumor growth after treatment with *S. aureus* microbial bioparticles indicating that our strategy was successful in inducing an effective anti-tumor response.

**Figure 2.**
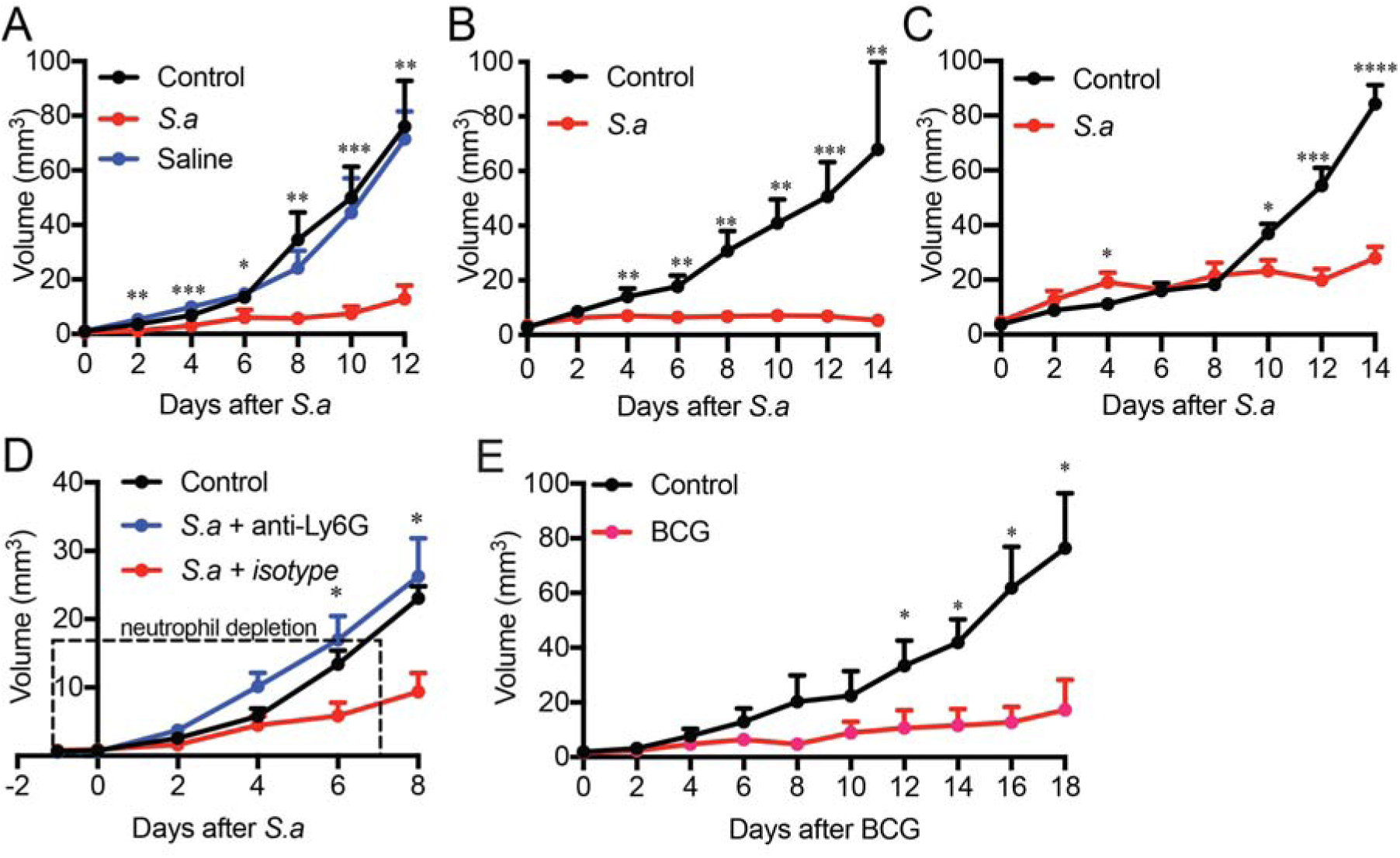
Microbial treatment leads to neutrophil-dependent inhibition of tumor growth. Tumor volume was measured in *S. aureus*, saline (vehicle) treated or control LLC (A), B16F10 3C (B) or AT3 (C) tumors grown in C57BL/6 mice and treated very second day. D. LLC tumor volume following anti-Ly6G neutrophil depletion on days −1, 1, 3, 5, 7. *S. aureus* was injected on days 0, 2, 4, 6, 8. E. Tumor volume was measured in BCG-treated or control LLC tumors in C57BL/6 mice. Data shown as mean + SEM of at least 3 tumors and are representative of 5 (A), 2 (B, D and E) and 1 (C) independent experiments. Data analyzed using Mann-Whitney test (A-B, D-E) or t-test (C) * P ≤ 0.05 ** P ≤ 0.01 *** P ≤ 0.001 **** P ≤ 0.0001.

To demonstrate that this is a broad phenomenon not restricted to a specific cancer model, we tested our microbial bioparticle treatment in two widely used pre-clinical tumor models, B16F10 melanoma and AT-3 breast cancer (**Fig. 2B and C**). In both models, microbial particle treatment substantially inhibited tumor growth. These results indicate that our approach is effective in a range of solid tumors.

### Neutrophils are required for tumor growth inhibition

To provide evidence of a causal contribution by MA neutrophils to the control of tumor growth in response to microbial treatment, we treated LLC-bearing mice with *S. aureus* bioparticles to suppress tumor growth but also depleted tumor-infiltrating neutrophils using neutrophil-specific anti-Ly6G antibody (**Suppl. Fig. 2**). Although neutrophils are hard to deplete long-term due to their rapid replenishment by the bone marrow, shortterm neutrophil depletion reversed the tumor suppression effect of *S. aureus* (**Fig. 2D**). These data demonstrate that MA neutrophils are essential for microbial bioparticle inhibition of tumor growth.

### Microbial tumor control is a broad phenomenon

We hypothesized that tumor growth inhibition achieved by treatment with *S. aureus* bioparticles is not unique to this bacterium and can be replicated using other microbes. To test this, we investigated whether administration of *Mycobacterium bovis* Bacillus Calmette Guerin (BCG) can modulate tumor growth. As was observed with *S. aureus* bioparticles, intratumoral treatment with BCG led to a rapid influx of neutrophils (**Suppl. Fig. 3**) and substantially inhibited tumor growth (**Fig. 2E**), demonstrating that the effect of microbial treatment on tumor growth is not restricted to particular microorganisms but is part of a broader phenomenon. This suggests that microbial agents like the BCG vaccine, which has a well-established clinical safety profile, can be repurposed for treating a range of solid tumors.

### Microbe- and tumor-derived signals shape neutrophil phenotype and turnover

Our data suggests that microbial bioparticle treatment may counteract the effects of the TME on neutrophils by converting them into anti-tumor effectors. However, little is known about the changes in neutrophil phenotype when subjected to signals within the TME. To investigate this, we took advantage of a photoconversion-based approach that we have developed to label tumor-infiltrating immune cells (17). Here we used it to label tumor infiltrating neutrophils and compare phenotypes of tumor-experienced neutrophils to neutrophils in circulation and neutrophils recently recruited to tumors. Our analysis showed that as neutrophils left circulation and infiltrated tumors they became activated (as indicated by increased expression of CD11b and downregulation in CD62L and CXCR2, **Fig. 3A-C**). Notably, this activation was more pronounced in photoconverted tumor-experienced neutrophils compared to non-photoconverted recently recruited neutrophils, indicating that circulating neutrophils undergo progressive activation upon entering tumors.

**Figure 3.**
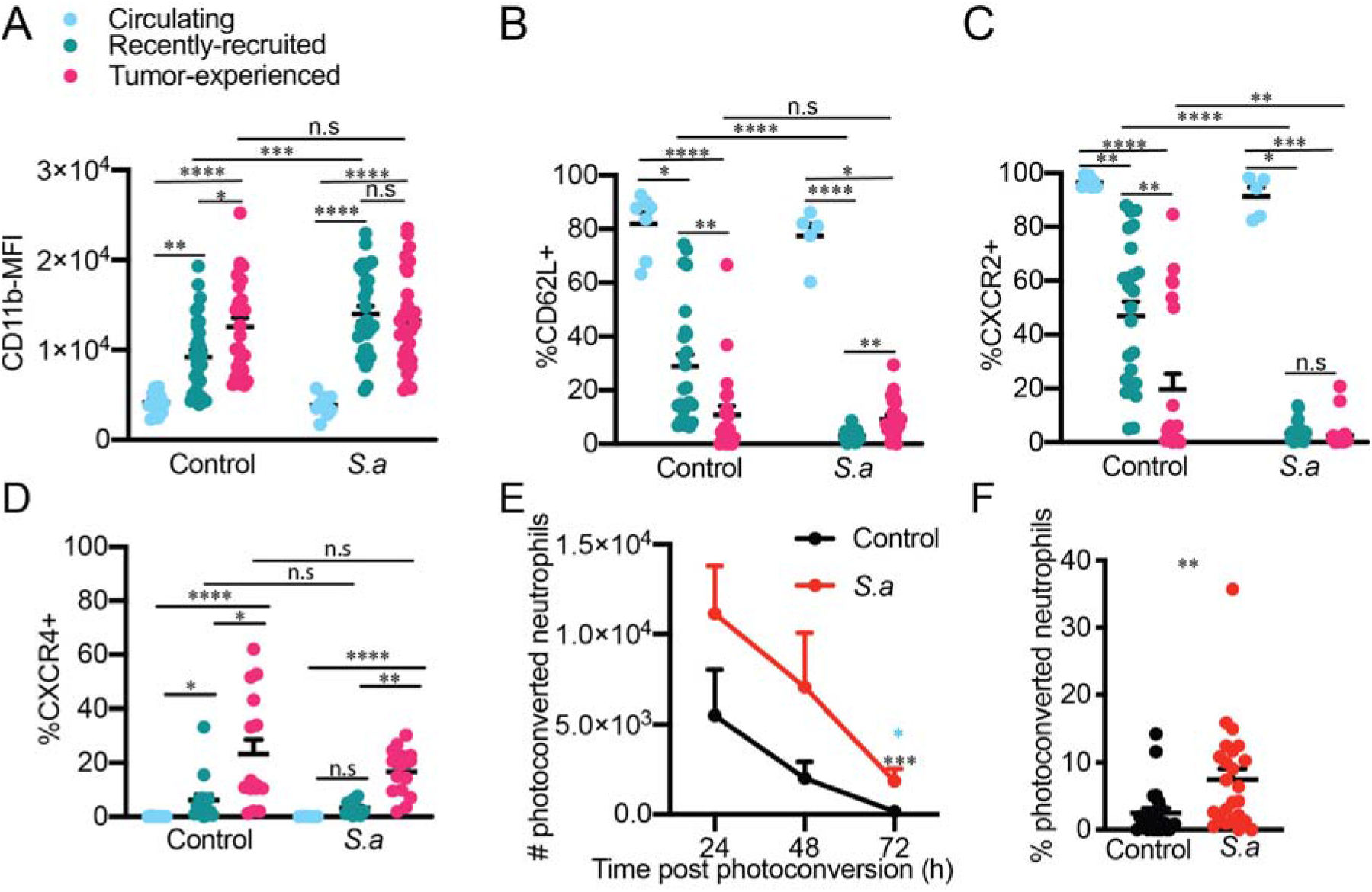
Neutrophil phenotype and persistence in tumors are modulated by signals within the TME. Tumors were photoconverted and treated with *S. aureus* bioparticles. CD11b (A), CD62L (B), CXCR2 (C) and CXCR4 (D) expression was analyzed 24 h later on circulating (blue), recently-recruited (non-photoconverted, green) and tumor-experienced (photoconverted, magenta) neutrophils using flow cytometry. E. Tumors were photoconverted and treated with *S. aureus* bioparticles immediately after photoconversion. Photoconverted neutrophils in tumors were quantified 24, 48 and 72 h later. F. Flow cytometric analysis of photoconverted neutrophils in tumor draining lymph 24 h after photoconversion. Data are pooled from two or more independent experiments. Each circle represents a tumor, blood or lymph node sample, mean +SEM are shown (A-D and F) or the mean+SEM of at least 7 tumor samples (E). Data was analyzed using one-way ANOVA (A), Kruskal-Wallis test (B-E), multiple T-test or Mann-Whitney test (F). * P ≤ 0.05 ** P ≤ 0.01 ***P ≤ 0.001 **** P ≤ 0.0001. In (E) Blue * indicates 24 vs 72 h comparision, black * indicates *S.aureus* vs control comparison.

When we examined the effect of microbial bioparticle treatment on phenotype of recently-recruited and tumor-experienced neutrophils, we found that neutrophil activation was enhanced by *S. aureus* bioparticles. However, tumor-experienced neutrophils were less susceptible to stimulation, suggesting that aged neutrophils lose some of their plasticity and ability to respond to signals in their environment.

We next used the photoconversion system to analyze the expression of chemokine receptor CXCR4, which is associated with neutrophil aging (18). As expected, CXCR4 expression was higher in photoconverted neutrophils that are likely to be more aged than non-photoconverted neutrophils that have recently entered the tumor from circulation (**Fig. 3D**). CXCR4 expression was unchanged in neutrophils from microbial particle-treated tumors suggesting that tumor neutrophils rapidly acquire an aging phenotype regardless of microbial signals.

Neutrophils are short-lived cells with a half-life of just several hours in circulation (19). However, their lifespan can be extended to around 2 days when neutrophils enter tissues (5). Despite neutrophils infiltrating most solid tumors, how long they survive in the TME is not yet known. To address this knowledge gap, we used photoconversion to label neutrophils in tumors and analyze the number of photoconverted neutrophils remaining in tumors at various timepoints. We observed a sharp decline in the number of photoconverted neutrophils in tumors over time – neutrophil number decreased by 63% between 24 and 48 hours and by 92% between 48 and 72 hours (**Fig. 3E**), suggesting that most tumor-infiltrating neutrophils do not survive for extended periods of time. The decline in photoconverted neutrophils was less rapid following *S. aureus* injection with neutrophil number decreasing by 37% between 24 and 48 hours and by 74% between 48 and 72 hours (**Fig. 3E**).

Neutrophil turnover in tumors is contributed to by the influx of neutrophils into tumors and death *in situ*. Whether neutrophils can also emigrate from tumors has not been examined. We applied photoconversion as previously (17) to assess neutrophil egress from tumors to draining lymph nodes. By photoconverting tumor-infiltrating cells prior to microbial bioparticle treatment and then analyzing draining lymph nodes for the presence of photoconverted tumor-egressing neutrophils, we found that neutrophils emigrated poorly from unmanipulated tumors (**Fig. 3F**). However, microbial bioparticle treatment substantially increased neutrophil egress to draining lymph nodes (**Fig. 3F**). Taken together our data indicate that microbial bioparticle treatment modulates neutrophil turnover within the TME and trafficking from tumors.

### Microbial treatment modulates neutrophil function in cancer

Increased neutrophil activation, coupled with neutrophil-mediated inhibition of tumor growth (**Figs. 1 and 2**), suggests that MA neutrophil functions are different from neutrophils in untreated tumors. To investigate this, we first examined uptake of bacterial particles by tumor neutrophils, since ingesting bacteria is a classical feature of neutrophil anti-microbial response. We observed that tumor-infiltrating neutrophils readily phagocytosed *S. aureus* (detected as uptake of labeled bioparticles) and were the main tumor immune subset to contain bacteria 24 hours following treatment (**Fig. 4A**). This demonstrates that microbial bioparticles elicit an anti-bacterial response in tumorinfiltrating neutrophils.

**Figure 4.**
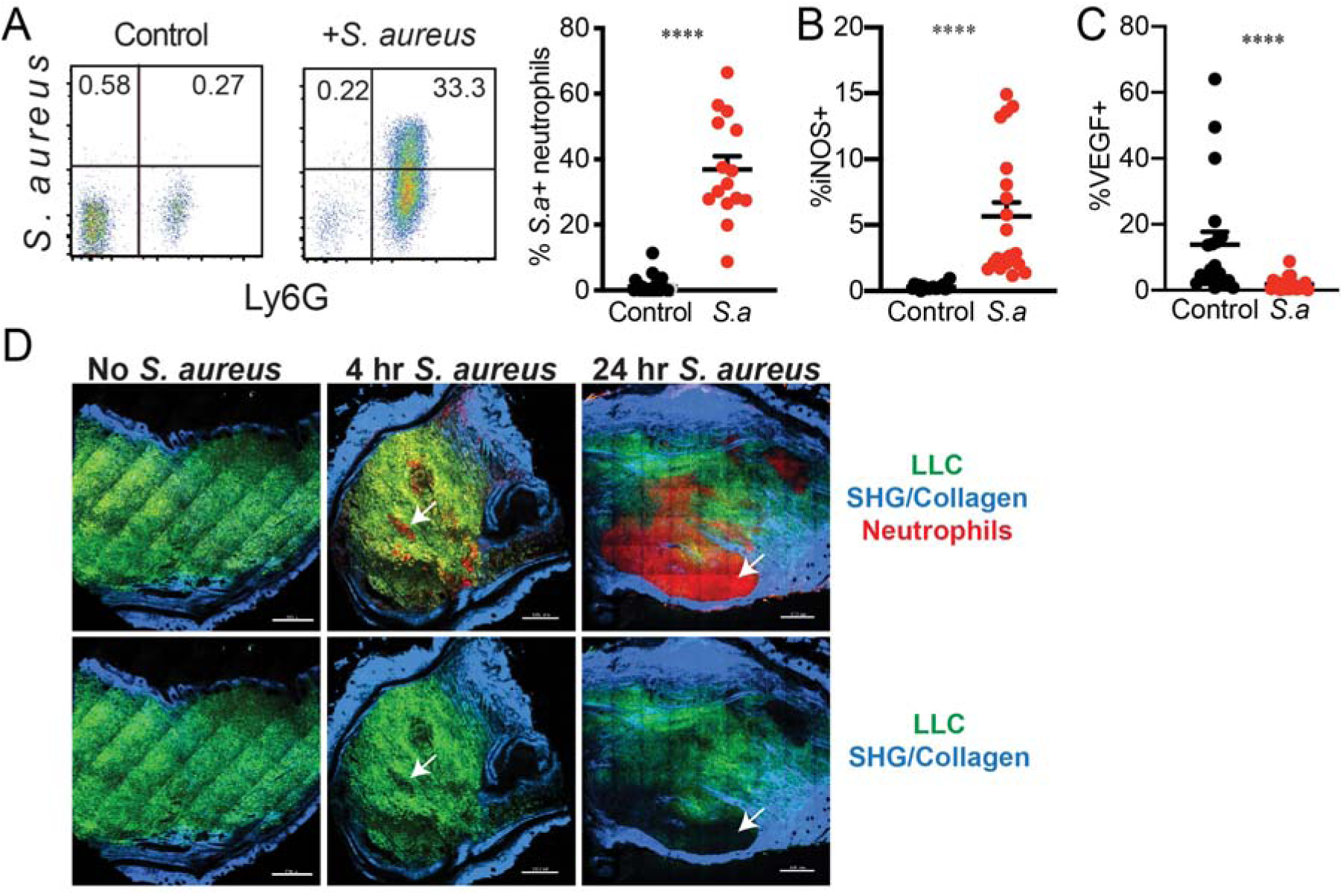
Change in tumor neutrophil function in response to microbial bioparticle treatment. A. Phagocytosis of labeled *S. aureus* bioparticles by intratumoral neutrophils. Representative flow cytometry plots (left) and pooled data (right) are shown. Proportion of iNOS (B) and VEGF (C) expressing neutrophils was quantiated by flow cytometry in control (black) and *S.aureus*-treated tumors (red). D. LLC-GFP tumors (green), neutrophils (red) and Second Harmonic Generation/Collagen (blue) were visualised in frozen sections from microbial bioparticle treated and control tumors using two-photon microscopy. White arrows indicate areas of tumor cell clearance by neutrophils. Scale bar represents 500μm. Data was pooled from three independent experiments and analyzed using Mann-Whitney test. Each circle represents a tumor, mean+SEM are shown. **** P ≤ 0.0001.

Phagocytosis induces extensive changes in neutrophil function (5). Therefore, we assessed neutrophil function in tumors before and after microbial treatment. We observed an increase in inducible nitric oxide synthase (iNOS) (**Fig. 4B**), an anti-microbial agent produced in response to phagocytosis that can also stimulate apoptosis and kill nearby cells by releasing reactive oxygen species (ROS) (20). We also noted a concomitant decrease in the production of Vascular Endothelial Growth Factor (VEGF) (**Fig. 4C**), one of the key pro-tumor molecules secreted by neutrophils in cancer (7). These changes in neutrophil function indicate that our microbial treatment drives a switch in neutrophil functional phenotype from pro-to anti-cancer effectors.

Increased production of iNOS following microbial treatment suggests that MA neutrophils may be remodeling surrounding matrix. To study neutrophil interactions with tumor cells we established LLC-eGFP tumors in the ear pinnae of neutrophil-specific reporter (BigRed/Catchup^IVM-red^) mice where tdTomato fluorescent protein is expressed in Ly6G^+^ neutrophils (21). When we examined tumor cross-sections 24 hours following microbial bioparticle treatment, we observed large neutrophil clusters within the tumor mass (**Fig. 4D**). Notably, these clusters corresponded to areas cleared of tumor cells and collagen suggesting that MA neutrophils in treated tumors may inhibit tumor growth by remodeling tumors and removing tumor cells.

### Migration of intratumoral neutrophils is controlled by signals in the TME

Despite their presence in most solid cancers, neutrophil dynamics in tumors have not been visualized. Our analysis indicated a change in neutrophil function following microbial treatment (**Fig. 4**). Since immune cell migration is tightly linked to their function, we hypothesized that the change in function will affect neutrophil dynamics in tumors. We leveraged intravital two-photon microscopy to visualize neutrophil dynamics within intact tumors in neutrophil reporter mice and identify how these change in response to microbial bioparticle treatment. Prior to treatment neutrophils were scattered throughout the tumor mass (**Fig. 5A**) and displayed limited motility (median average speed 0.04 μm/second) and displacement (median displacement 3 μm) (**Fig. 5B and C and Supplementary Video 1**).

**Figure 5.**
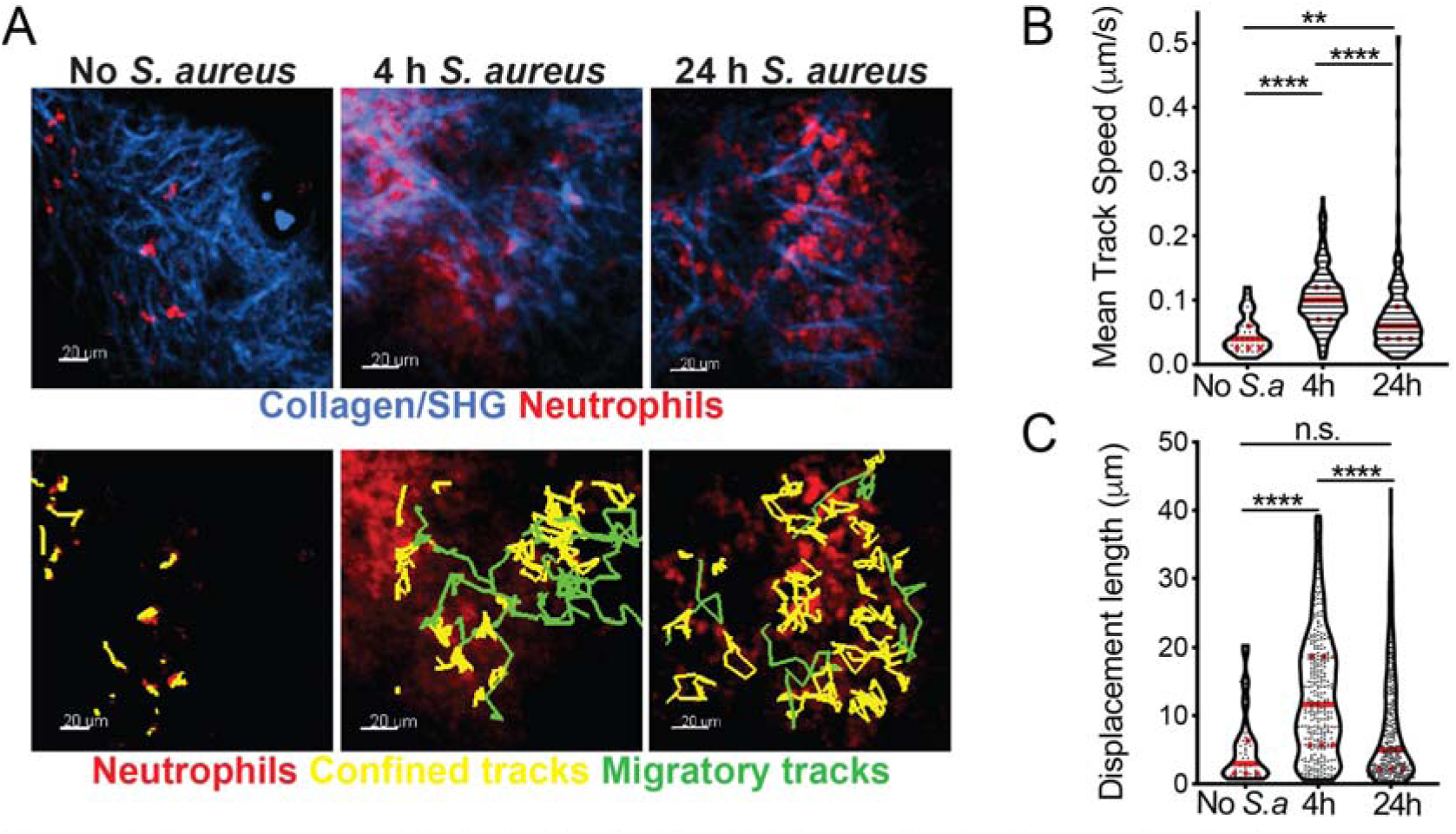
Tumor neutrophil dynimics *in vivo*. A. Neutrophils (red) were visualized in steady state or in *S. aureus* (*S.a*) bioparticle treated LLC tumors using intravital two-photon microscopy. Yellow tracks indicate neutrophils with confined motility (track displacement length<l3μm), green tracks indicate migrating neutrophils (track displacement length>47 μm). Second Harmonic Generation(SHG)/colla-gen - blue. Bar represents 20 μm. B. Mean track speed of intratumoral neutrophils. C. Track displacement length of intratumoral neutrophils. Data from at least 4 independent imaging experiments per time point was analysed using a one-way ANOVA with Dunn’s correction for multiple comparisons (B and C). Median and quartiles are shown. **P ≤ 0.01, **** P ≤ 0.0001, n.s. not significant.

However, dynamics and distribution of neutrophils rapidly changed following microbial treatment. As early as 4 hours after treatment, we detected large clusters of neutrophils throughout the tumor mass (**Fig. 5A**). We also observed a 2.5-fold increase in neutrophil speed and a 4-fold increase in displacement (**Fig. 5B and C and Supplementary Video 1)**. The rise in motility was transient and by 24 hours after microbial treatment, neutrophil speed started to decline, although was still elevated in comparison to unmanipulated tumors.

Detailed analysis of MA neutrophil motility post treatment revealed two distinct modes of behavior, where a proportion of neutrophils remained motile, while most neutrophils formed large stable clusters of low motility (**Fig. 5A**, **Supplementary Video 1**). These results show that microbial bioparticle treatment altered neutrophil dynamics in tumors – MA neutrophils displayed increased motility and formed large clusters within the tumor mass.

### MA neutrophils mediate tumor cell destruction

Neutrophil depletion experiments (**Fig. 2D**) provided evidence that neutrophils are important for tumor growth control, while analysis of tumor sections (**Fig. 4D**) suggested that this may be mediated by neutrophil remodeling of tumors and removal of tumor cells. To test this, we applied intravital imaging to study interactions between neutrophils and LLC-eGFP tumor cells *in vivo* in real time in neutrophil reporter mice. Using this approach, we detected an increase in rounded tumor cells and tumor cell debris in areas of neutrophil infiltration in treated tumors, in contrast to elongated intact LLC cells in unmanipulated tumors (**Fig. 6A, Supplementary Video 2**). Quantitation of eGFP signal showed a loss of LLC tumor cells (**Fig. 6A)**, while labeling with SYTOX confirmed tumor cell death within neutrophil clusters (**Fig. 6B)** in treated tumors.

**Figure 6.**
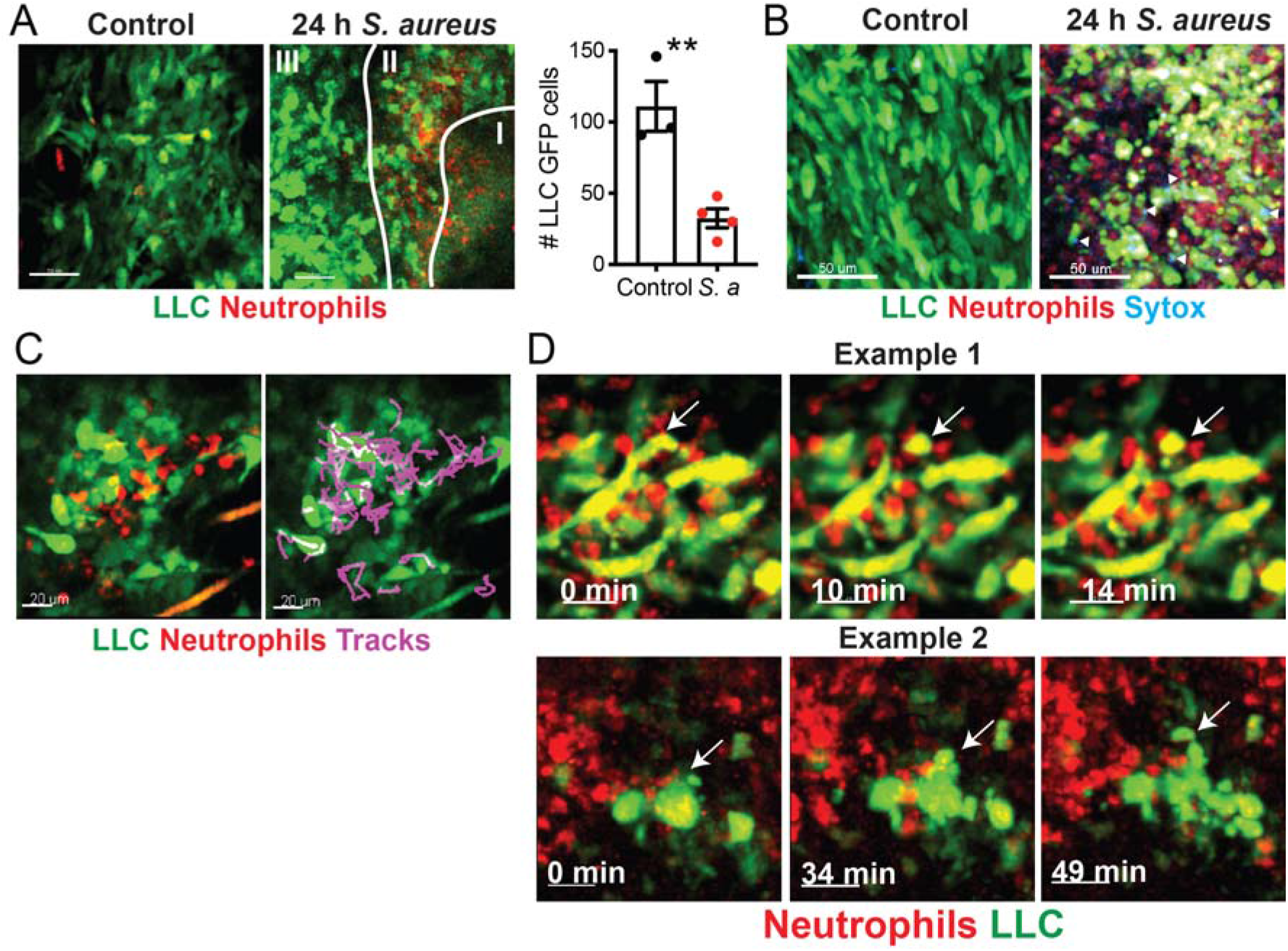
Neutrophil-tumor cell interactions *in vivo*. A. Neutrophils (red) and LLC cells (green) were visualized in tumors prior or 24h following *S. aureus* bioparticle treatment using intravital two-photon microscopy. Region 1 indicates an area of LLC loss, region II indicates partial LLC destruction, region III indicates an area of largely intact but rounded LLCs. Loss of LLC-GFP cells was quanitated (right panel, mean +/-SEM). B. LLC-GFP tumors (green), neutrophils (red) and SYTOX (blue) were visualised in frozen sections from microbial bioparticle treated and control tumors using two-photon microscopy. White arrows indicate SYTOX staining. Scale bar represents 5Oμm. C. Purple tracks show neutrophils (red) interacting and crawling over LLC (green). Bar represents 20 μm. D. Two examples of neutrophil-LLC interactions and tumor cell blebbing (white arrows). Bar represents 20 μm. **p ≤ 0.01.

When we visualized neutrophil-tumor cell interactions over time, we observed that neutrophils engaged in multiple interactions with tumor cells (**Fig. 6C, Supplementary Video 3**). Detailed analysis of these interactions revealed tumor cells within neutrophil clusters undergoing blebbing suggesting that neutrophils remove tumor cells *in vivo* (**Fig. 6 D**, **Supplementary Video 4)**. Taken together our results indicate that MA neutrophils mediate microbe-induced tumor growth control by destroying tumor cells.

## DISCUSSION

Immunotherapy, as demonstrated by the success of drugs disrupting immune checkpoints, can unleash anti-tumor immunity and mediate durable cancer regression. Here we have expanded upon this concept to develop a new cellular target for anti-cancer immune therapy, namely MA neutrophils which can provide a strong anti-cancer effect. These findings are in contrast with the prevailing view of neutrophils in cancer as pro-tumorigenic cells. Consistent with a tumor-promoting role for neutrophils, depleting this immune subset has been demonstrated to be an effective strategy in inhibiting tumor growth and metastasis in several mouse models of cancer (12,14,22). However, because of their indispensable role in defense against bacteria, depleting neutrophils can expose patients to severe and life-threatening infection (23). Furthermore, neutrophils are hard to deplete for a substantial period of time as they are rapidly regenerated by the bone marrow. Instead of depleting neutrophils, our approach takes advantage of their natural anti-microbial capabilities and redirects this activity against tumor cells. Microbial bioparticle treatment recruited, activated and polarized neutrophils in tumors to generate MA neutrophils with a capacity to destroy tumor cells. Microbial bioparticle treatment induced a switch in neutrophil function from VEGF production, which stimulates angiogenesis and favors tumor growth (7), to increase in iNOS, which may contribute to tissue remodeling and tumor cell death. MA neutrophils successfully inhibited tumor growth in three pre-clinical tumor models indicating that neutrophil plasticity in cancer could be exploited across a range of solid tumors.

We applied two-photon microscopy to uncover unique MA neutrophil dynamics, phenotype and functions that are shaped by opposing signals within the TME. In the unmanipulated TME, intratumoral neutrophils show limited displacement and motility. This is in contrast with the rapidly migrating neutrophils that respond to infection by coalescing together to form dynamic swarms (24). Since immune cell migration reflects their function, it is likely that limited intratumoral neutrophil migration indicates a distinct pro-tumor phenotype induced by signals within the TME (25).

Microbial bioparticle treatment led to rapid influx and activation of neutrophils in tumors, inducing an intratumoral neutrophil phenotype similar to that of neutrophils responding to infection in healthy tissues (26). Consistent with the change in surface phenotype, treatment with microbial bioparticles led to a substantial increase in intratumoral neutrophil motility indicating a change in neutrophil function.

In infection and sterile injury neutrophil swarms remodel underlying tissue and remove collagen (24,27). This process is thought to be mediated by release of ROS and matrix remodeling enzymes (28). Microbial bioparticle treatment led to formation of neutrophil clusters, that corresponded to areas of tumor cell clearance, and a concomitant increase in production of iNOS, which is associated with release of ROS and tissue remodeling (5). Real time intravital imaging in tumors revealed that neutrophil interactions with cancer cells led to tumor cell blebbing indicating that neutrophils can directly destroy tumor cells *in vivo*. Notably, we did not observe transfer of cancer cell plasma membrane to neutrophils suggesting that, unlike ADCC-mediated cancer cell death (29), tumor cell death did not occur by neutrophil trogocytosis.

Neutrophils are exquisitely sensitive to signals in their microenvironment (30). Our microbial particle treatment led neutrophils to assume a highly activated state. However, this effect was transient and activation subsided to pre-stimulation levels within three days. This suggests a dynamic balance between opposing signals provided by microbial bioparticles and the TME - as microbial signals diminish, neutrophils can revert to a tumor-induced phenotype. This plasticity of intratumoral neutrophils is an important consideration for treatments that lead to neutrophil recruitment and activation such as surgery and radiation. Microbial bioparticle treatment could provide benefit as adjuvant therapy together with surgery and radiation to ensure that recruited neutrophils take on an anti-tumor phenotype.

Although there are important differences between mouse and human neutrophils, studies dating as far back as 1891, when bone surgeon William Coley observed that an antimicrobial response may be beneficial in solid cancers (31), suggest that bacteria can effectively stimulate tumor immunity in humans. We observed a strong anti-tumor response using not only killed *S. aureus* bioparticles but also BCG, which is used worldwide as a vaccine against tuberculosis and post-surgery to prevent relapse in bladder cancer but so far has limited applications in other solid cancers (32). This suggests that bacterial vaccines with proven clinical safety profiles can be repurposed for treating a range of solid tumors, paving the way for broader utilization of microbial immunotherapy in cancer (33,34). However, our study highlights the need for understanding neutrophils activation, polarization and turn-over in tumors in order to optimize bacterial-based therapy as a treatment for cancer.

Immunosuppression mediated by cancer cells is a key obstacle for an effective anti-tumor response. Our microbial bioparticle treatment drastically reshapes the TME by recruiting a large number of MA neutrophils with anti-rather than pro-tumor functions. Furthermore, neutrophils are emerging not only as a critical aspect of host defense against bacteria but also as a controller of adaptive immunity (5,26). Therefore, recruiting MA neutrophils may provide a pathway to increase adaptive immune infiltration and activation in poorly infiltrated cancers (e.g. breast or pancreatic cancers), where insufficient immune infiltration and immunosuppression have been major obstacles for checkpoint inhibitor therapy. This suggests that combination strategies simultaneously targeting both innate and adaptive immune systems may synergize to overcome checkpoint blockade resistance and promote tumor killing.

## MATERIAL AND METHODS

### Mice

All mice used in this study were maintained on C57BL/6 (RRID:MGI:5656552) background and housed in specific-pathogen free conditions. All animal experiments and procedures were approved by the Garvan Institute of Medical Research/St Vincent’s Hospital Animal Ethics Committee. Male and female tumor bearing mice were randomly assigned to treatment groups once their tumors were established. C57BL/6 mice (RRID:MGI:5656552) were obtained from Australian BioResources (Moss Vale, NSW). Kaede mice (35) were a gift from Professor Michio Tomura and were backcrossed and maintained on the C57BL/6 background. Ly6G Cre tdTomato (C57BL/6-*Ly6g*(tm2621(Cre-tdTomato)Arte) neutrophil-specific reporter mice (21) were a gift from Professor Matthias Gunzer and crossed with B6.LSL td-Tomato (B6.Cg-*Gt(ROSA)26Sor^tm14(CAG-tdTomato)Hze^*/J (IMSR Cat# JAX:007914 RRID:IMSR_JAX:007914) to generate BigRed/Catchup^IVM-red^ mice (36).

### Tumor cell lines

Mouse Lewis Lung Carcinoma (LLC) (ATCC Cat# CRL-1642, RRID:CVCL_4358) cell line was purchased from ATCC. LLC-eGFP cells were a gift from Professor Robert Brink. B16F10-3C melanoma cell line was a gift from Professor Wolfgang Weninger. AT-3 (RRID:CVCL_VR89) breast cancer cell line was a gift from Dr Scott Abrams.

### Microbial bioparticles and microbes

*S. aureus* bioparticles (Wood strain without protein A) (Thermofisher) were resuspended in PBS with 2 mM sodium azide and 4–20×10^6^ bioparticles were injected directly into tumors. *Mycobacterium bovis* Bacillus Calmette Guerin were prepared as described previously (37) and 5×10^6^ CFU were injected directly into tumors every second day.

### Neutrophil recruitment into tumors

LLC cells were inoculated into ear pinnae of Kaede mice. When tumors reached 4-8 mm^3^, they were photoconverted for 20 minutes with a violet light from a cold-light source fitted with a filter (Zeiss) to minimize thermal and phototoxicity (17,26) and immediately injected with 20×10^6^ *S. aureus* bioparticles. Twenty-four, 48 and 72 h later mice were sacrificed and tumors, draining lymph nodes and blood were analyzed by flow cytometric analysis.

### Microbial control of tumor growth

LLC, B16F10-3C, AT-3 tumor cells were inoculated into ear pinnae of C57BL/6 mice. Once tumors were detected, they were treated with 4 – 20×10^6^ *S. aureus* bioparticles or 5×10^6^ CFU BCG or saline vehicle administered intratumorally every 2 days until mice reached ethical endpoints. Tumor dimensions were measured using calipers and volume calculated with the modified ellipsoidal formula (V = ½ (Length × Width^2^).

### Neutrophil depletion

LLC tumors were grown in the ear pinnae of C57BL/6 mice. Once tumors became visible, neutrophils were depleted by intraperitoneal injection of 500 μg anti-Ly6G clone 1A8 (Bio X Cell Cat# BE0075-1, RRID:AB_1107721) or rat IgG2a isotype control clone 2A3 (Bio X Cell Cat# BE0089, RRID:AB_1107769). Twenty-four hours later 4-10×10^6^ *S. aureus* bioparticles were injected into LLC tumors. The mice then had alternating days of maintenance dose of 250 μg anti-Ly6G/isotype i.p. and *S. aureus* bioparticles injected into tumors every 2 days. A total of 4 doses of 250 μg anti-Ly6G or isotype was administered whilst *S. aureus* bioparticles were given until mice reached ethical endpoints.

Neutrophil depletion in mice treated with anti-Ly6G was confirmed by flow cytometric analysis. Blood samples red cells were lysed with 10 mM KHCO_3_, 0.1 mM EDTA and 166 mM NH4Cl solution and then blocked with 5% normal mouse serum and then stained with unlabeled rat anti-mouse-Ly6G clone 1A8 (Bio X Cell Cat# BE0075-1, RRID:AB_1107721) primary antibody, washed with FACS buffer and then stained with a secondary goat anti-rat IgG DyLight 649 (BioLegend Cat# 405411, RRID:AB_1575141) antibody and then washed with FACS buffer and blocked with 5% normal rat serum. Samples washed again with FACS buffer and then stained with labeled cell surface antibodies including CD11b-APCef780 clone M1/70 (Thermo Fisher Scientific Cat# 47-0112-80, RRID:AB_1603195).

### Flow cytometry and antibodies

Single cell suspensions were made from tumors and lymph nodes by mechanical disruption and passed through 100 μM strainers. For blood samples red cells were lysed. Staining for flow cytometry was performed in 96 well plates. Cells were blocked with CD16/CD32 clone 93 (Thermo Fisher Scientific Cat# 14-0161-86, RRID:AB_467135) for 15 min and stained with surface antibodies on ice in FACS buffer (1× PBS + 0.2% BSA and 0.1% NaN3 + 2 mM EDTA) for 30 min in the dark.

Cells for intracellular cytoplasmic staining were fixed with IC Fixation Buffer (Thermofisher) for 30 min following surface staining. Samples were washed twice in Permeabilization Buffer (Thermofisher) and stained in the same buffer with intracellular antibodies for 30 min at room temperature. Cells were washed twice in Permeabilization Buffer and resuspended in FACS buffer prior to flow cytometric acquisition. All samples were acquired on LSRII flow cytometer (BD Bioscience). Data was analyzed using FlowJo (FlowJo, RRID:SCR_008520).

### Two-photon intravital microscopy

For neutrophil motility imaging experiments LLC cells were inoculated into the ear pinnae of BigRed/Catchup^IVM-red^ mice. Tumors were imaged ~10-14 days later. For treated tumors, *S. aureus* 20×10^6^ bioparticles were injected into tumor and 4 – 24 h later tumors were imaged using two-photon microscopy. LLC-eGFP cells were used to visualize LLC cells inside tumors in order to analyse interactions between neutrophils and tumor cells.

Intravital two-photon microscopy was based on a previously described method (38). Two-photon imaging was performed using an upright Zeiss 7MP two-photon microscope (Carl Zeiss) with a W Plan-Apochromat 20×/1.0 DIC (UV) Vis-IR water immersion objective. Four external non-descanned detectors were used to detect blue (SP 485), green (BP 500-550), red (BP 565-610) and far red (BP 640-710). High repetition rate femtosecond pulsed excitation was provided by a Chameleon Vision II Ti:Sa laser (Coherent Scientific) with 690-1064nm tuning range. We acquired 3μm z-steps at 512×512 pixels and resolution 0.83μm/pixel at a frame rate of 10 fps and dwell time of 1.27 μs/pixel using bidirectional scanning. Anesthesia was induced with 100mg/kg ketamine/5mg/kg xylazine and maintained with 1-2% isoflurane supplemented with 100% oxygen at a flow rate of 500ml/min via a nose cone. Anesthetized mice were kept warm using a customized heated SmartStage (Biotherm). The ear was immobilized on a base of thermal conductive T-putty (Thermagon Inc.) using Vetbond tissue adhesive (3M).

### Image processing and data analysis

Raw image files were processed using Imaris (Imaris, RRID:SCR_007370) software. A Gaussian filter was applied to reduce background noise. Tracking was performed using Imaris spot detection function to locate the centroid of cells. Motility parameters such as cell displacement (or track length calculated as the total length of displacements within the track) and track speed (calculated by dividing track length by time) were obtained using Imaris Statistics function. All modelling and statistical analysis was performed in GraphPad Prism (GraphPad Prism, RRID:SCR_002798).

To quantitate LLC-eGFP cells, Imaris spot detection function was used to identify eGFP cells in the green channel for image areas of 250×250 μm in untreated and bioparticle treated tumors. The number of LLC cells remaining in each area was then calculated and recorded.

### Whole tumor section microscopy

Tumors were harvested from BigRed/Catchup^IVM-red^ mice and fixed for 45-90 min in the dark at room temperature in fixing buffer (1×PBS, 4% Formalin and 10% Sucrose). Subsequently, tumors were sequentially incubated with 10%, 20% and 30% Sucrose in PBS for 8 h in the dark at 4°C, embedded in O.C.T. compound (Sakura Finetek) and frozen at −80 °C. One hundred μm-thick cryosections of LLC tumor were cut and imaged using two-photon microscopy. Acquired images were processed and analyzed on Imaris (Bitplane) software.

### Statistical analysis

The statistical distribution of experimental data was determined using a D’Agostino-Pearson omnibus normality test. The statistical significance of experimental data for comparisons of two groups was determined using either an unpaired T-test when distribution was normal and unpaired Mann-Whitney test if distribution was not normal. Statistical significance, for comparison of three or more groups unmatched ordinary oneway ANOVA test when distribution was normal and unmatched Kruskal-Wallis test if distribution was not normal. All analysis was done on GraphPad Prism (GraphPad Prism, RRID:SCR_002798).

**Table 1.**
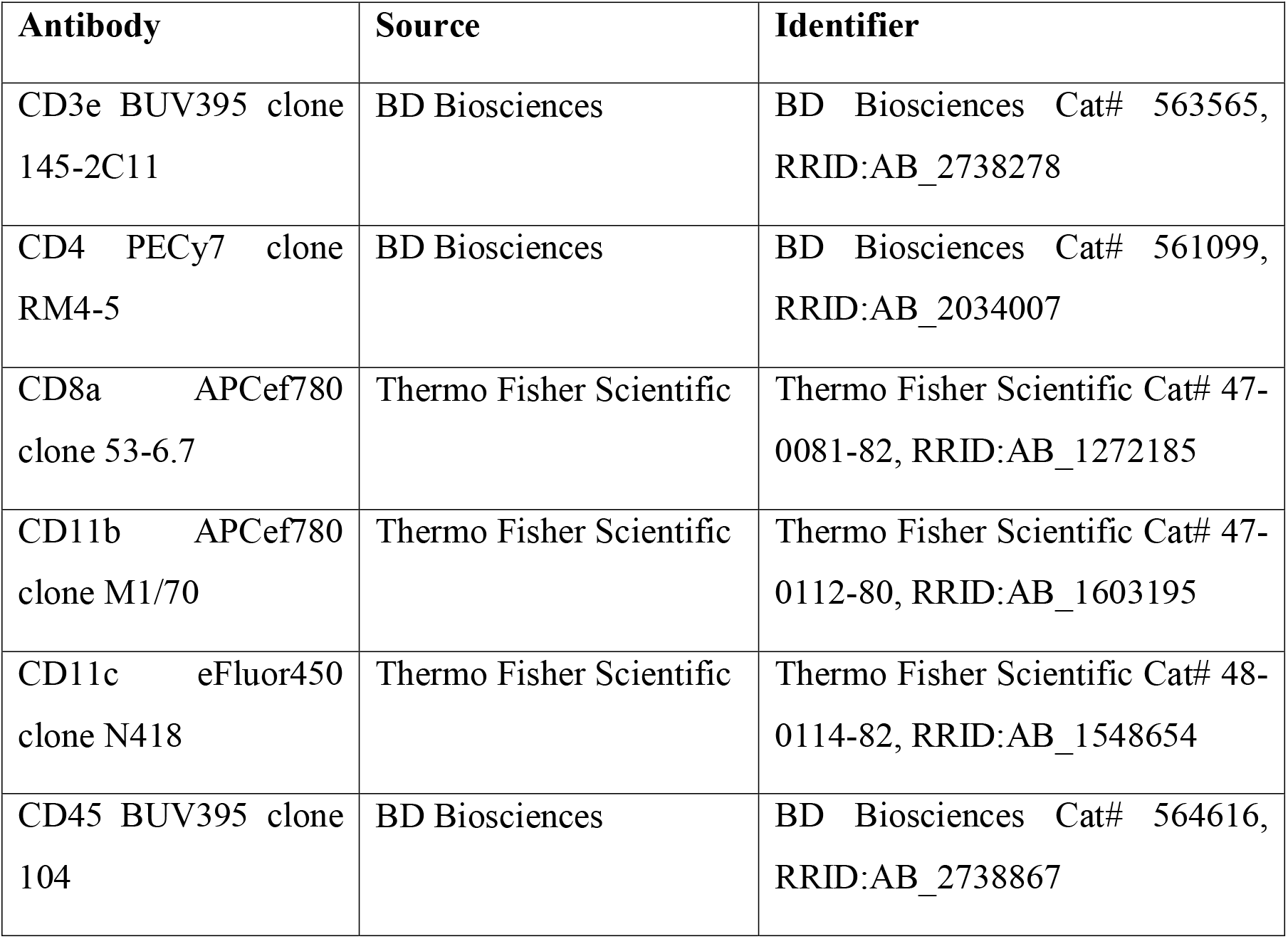
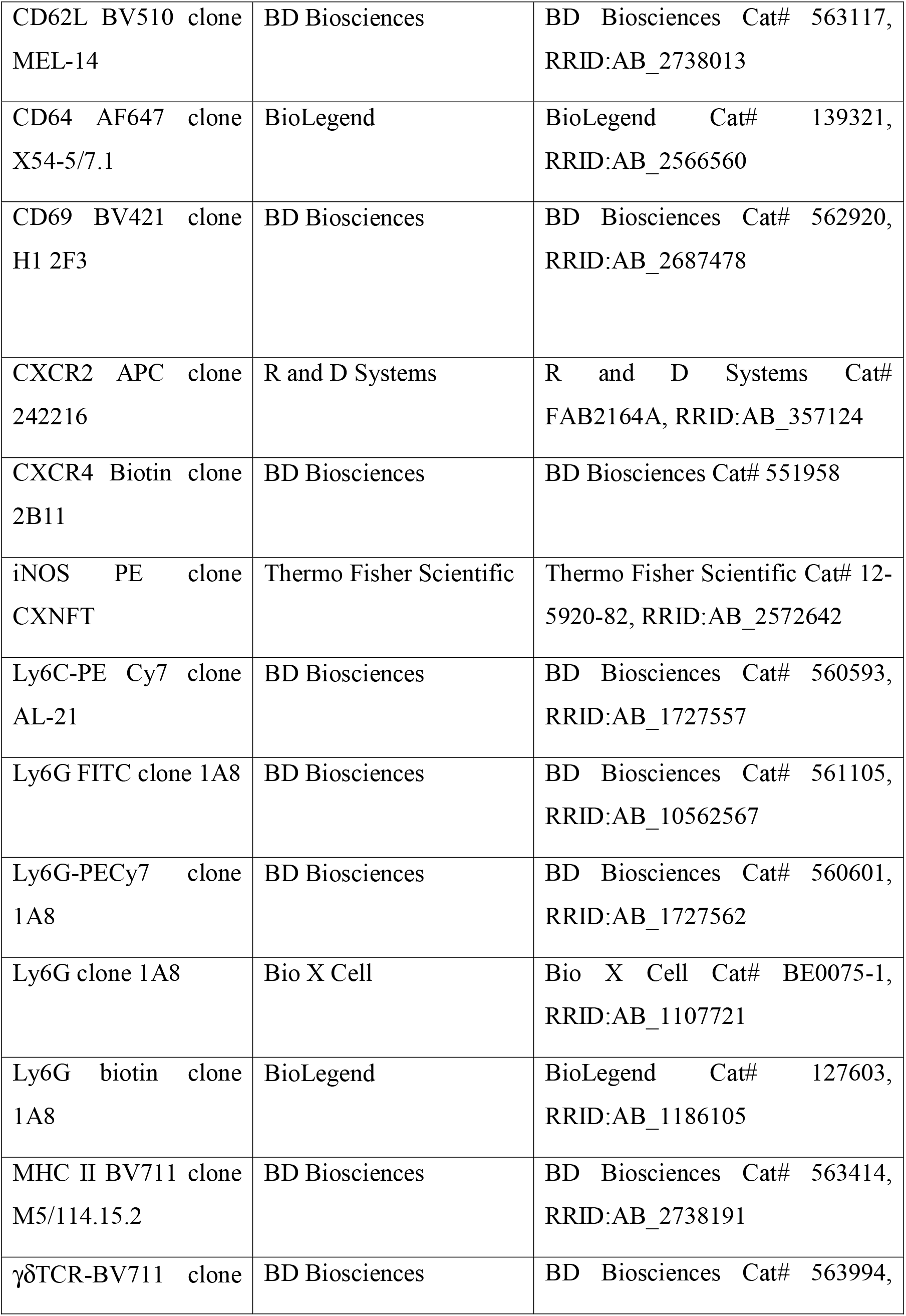
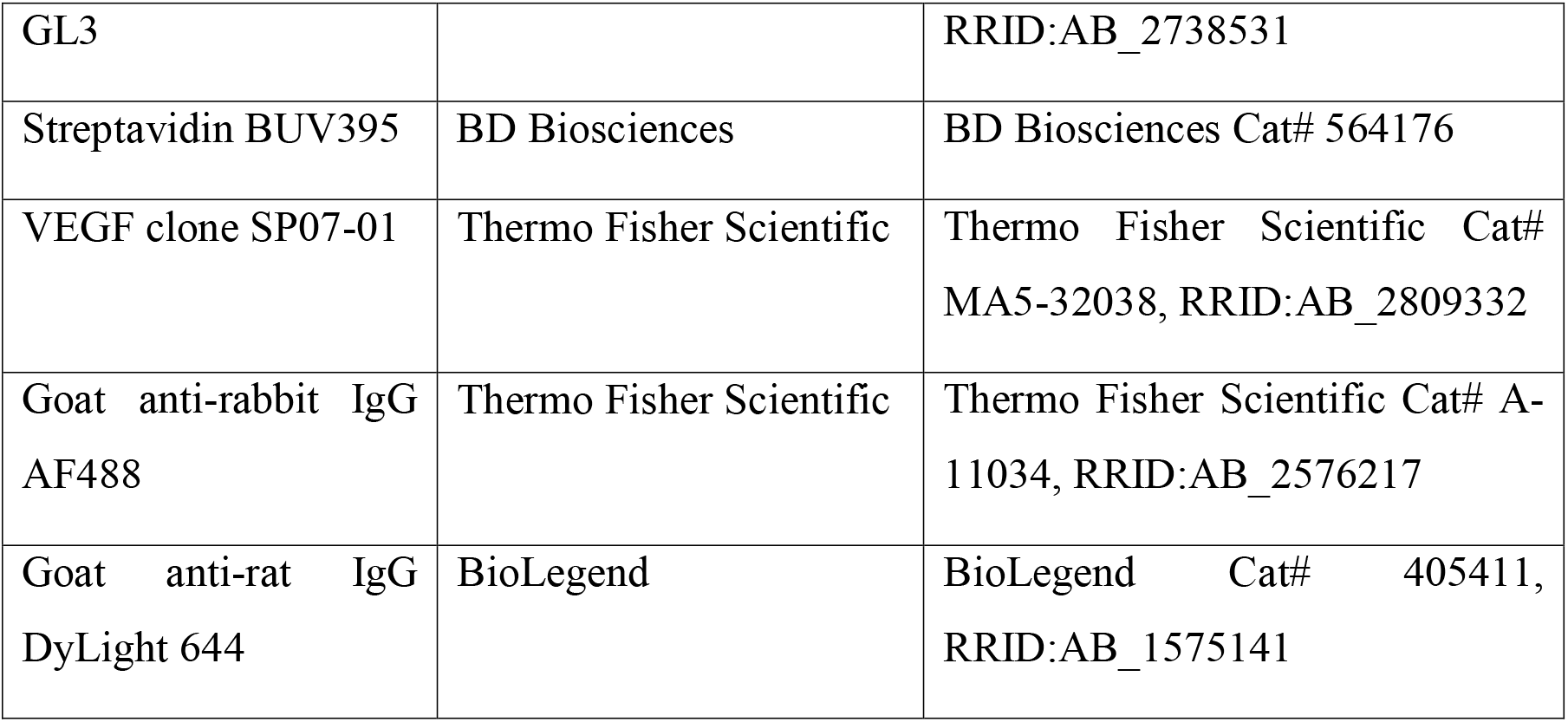
Antibodies for flow cytometry

## Notes

**Funding support:** This research was funded by National Health and Medical Research Council project grant GNT1106043, National Breast Cancer Foundation IIRS-19-027 (T.C.), Australian Government Research Training Program and Royal Australasian College of Physicians Fellows Research Entry Scholarships, Phil Salter Immuno-Oncology Fellowship (A.O.Y.) and supported by Peter and Val Duncan.

The authors declare no potential conflicts of interest.

### Competing Interest Statement

The authors have declared no competing interest.

