## Supplementary figures and images for "Microbial activation converts neutrophils into anti-tumor effectors"

### Supplementary Figures

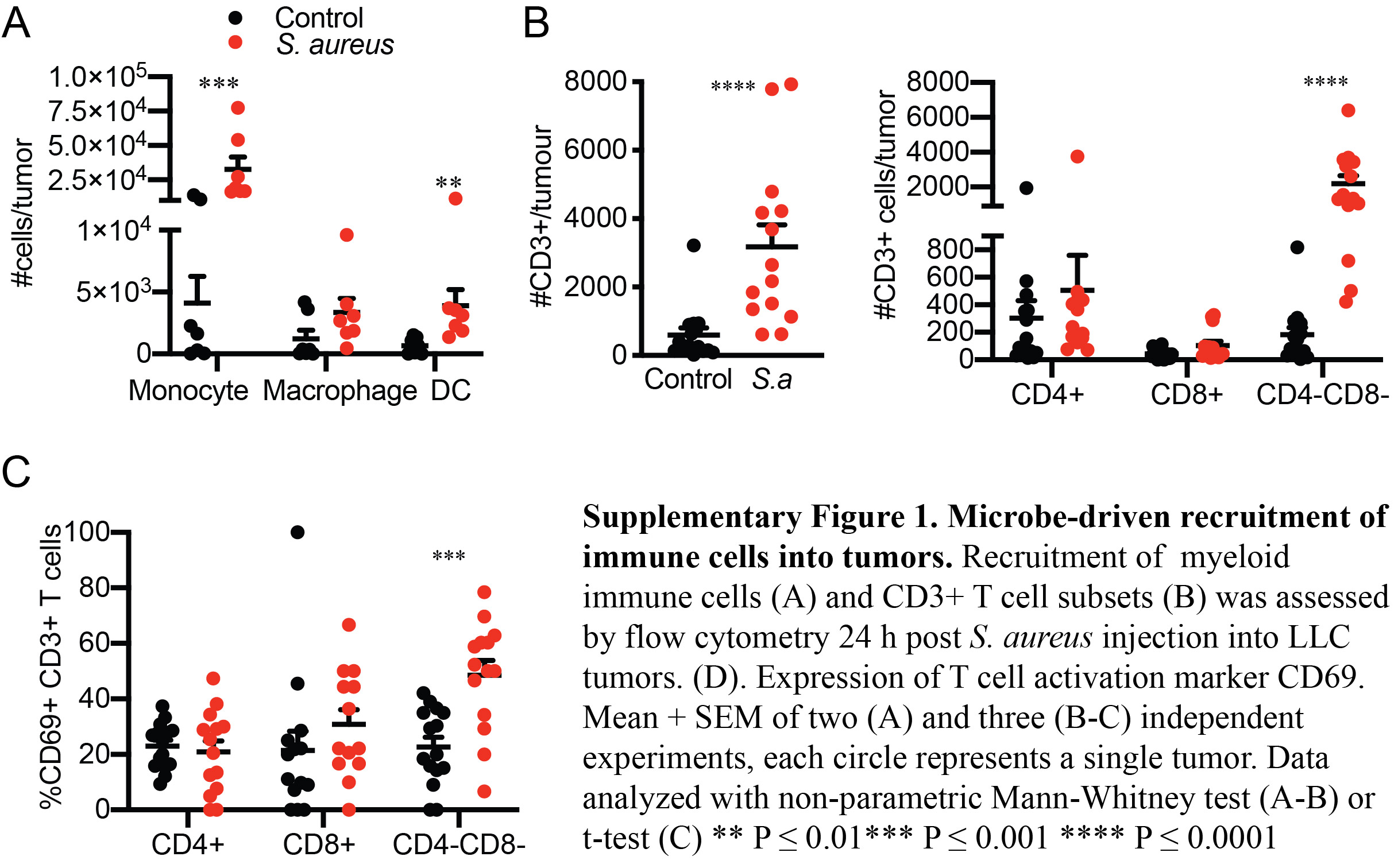

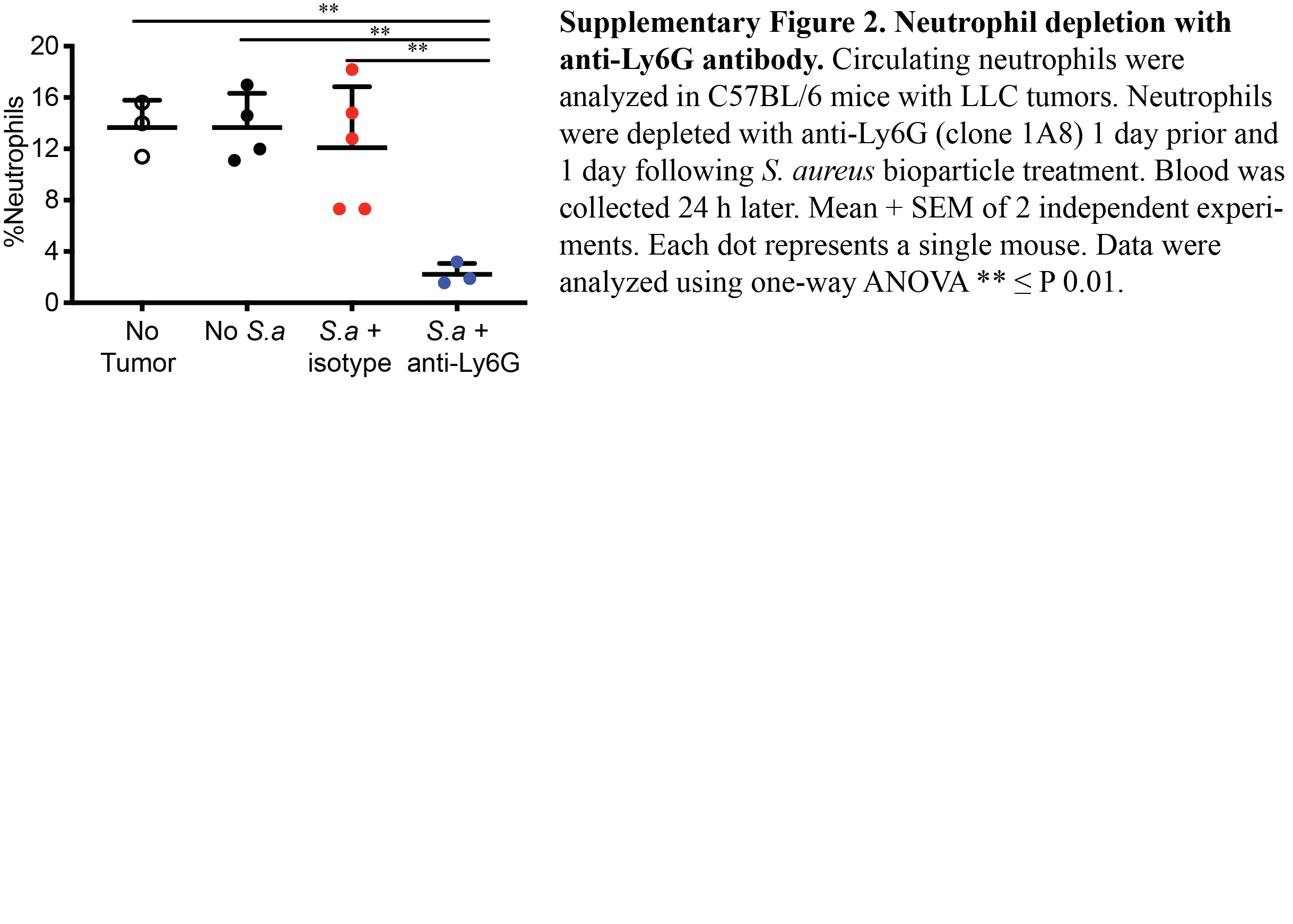

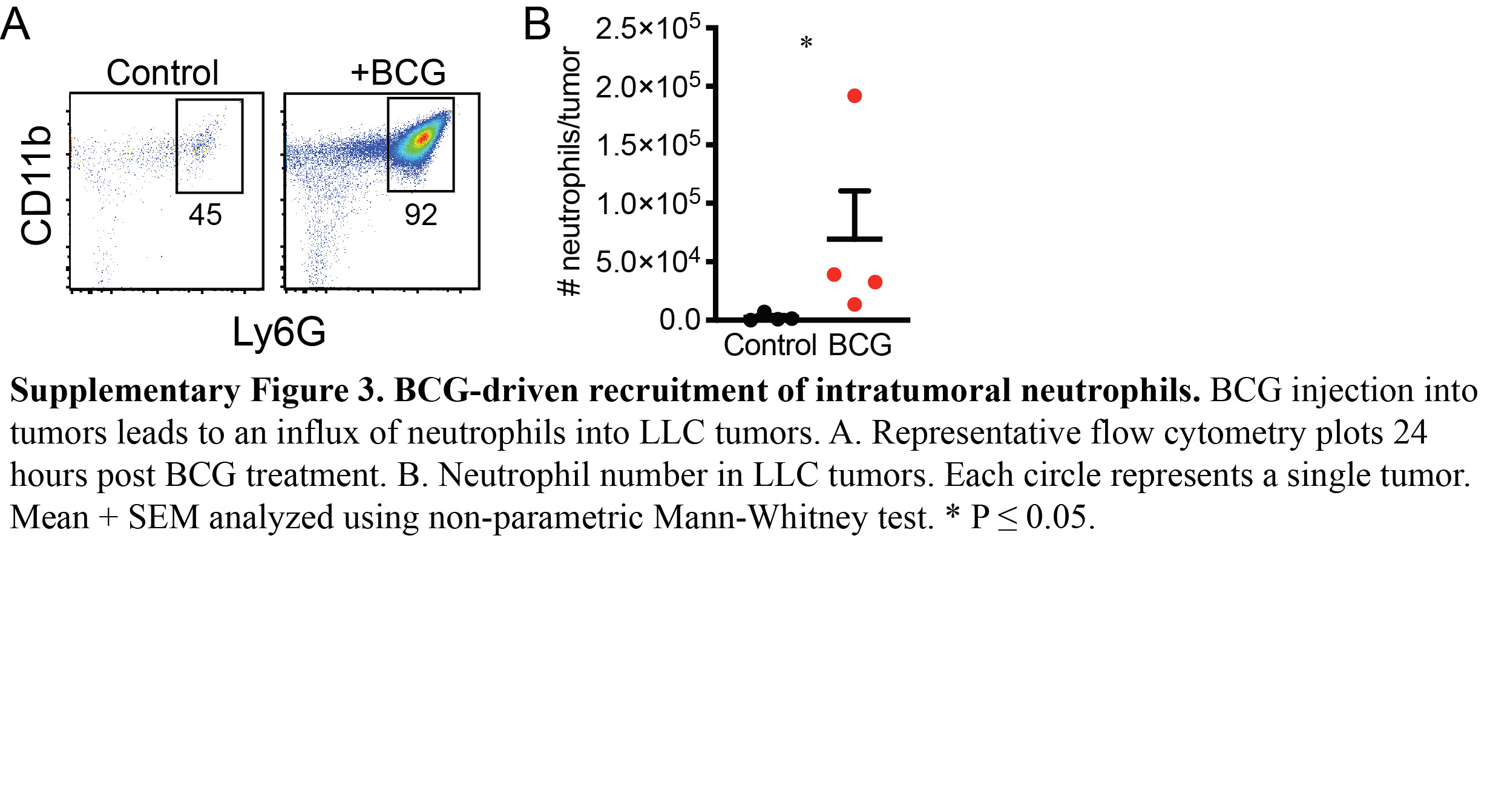
