## Supplementary material for "Microbial activation converts neutrophils into anti-tumor effectors": Video legends

**Supplementary VIDEO LEGENDS**

**Video 1. Neutrophil migration before and after microbial bioparticle treatment.** Ly6G neutrophil (red) migration in intact unmanipulated LLC tumors (part 1) and tumors that were treated with *S. aureus* bioparticles 4 and 24 hours earlier (part 2 and 3 respectively). Tracks indicate neutrophil paths. Yellow tracks indicate neutrophils with restricted motility (track displacement length less than 13 μm), green tracks indicate more motile neutrophils (track displacement length greater than 47 μm). Collagen/Second harmonic generation (SHG, blue). Projection of a time-lapse series imaged in LLC tumors in the ear flap of BigRed/Catchup^IVM-red^ mice. Part 1 dimensions: 213 μm x 213 μm x 45 μm x 20 minutes. Images were taken 30 seconds apart; Part 2 dimensions: 166 μm x 166 μm x 93 μm x 30 minutes. Images were taken 30 seconds apart; Part 3 dimensions: 191 μm x 191 μm x 24 μm x 48 minutes. Images were taken 60 seconds apart. Elapsed time is shown as days:hours:min:seconds.

**Video 2. Neutrophil interactions with LLC cells in unmanipulated and treated tumors.** Part 1: Intact elongated LLC (green) cells in unmanipulated tumors with scattered Ly6G neutrophils (red). Part 2: LLC cells progressively lose their elongated appearance to become rounded and then be broken down by Ly6G neutrophils. Projection of a time-lapse series imaged in eGFP-LLC tumors in the ear flap of BigRed/Catchup^IVM-red^ mice. Part 1 dimensions: 249 μm x 249 μm x 105 μm x 20 minutes. Images were taken 30 seconds apart; Part 2 dimensions: 249 μm x 249 μm x 105 μm x 49 minutes. Images were taken every 60 seconds. Elapsed time is shown as days:hours:min:seconds.

**Video 3. Neutrophils engage in multiple interactions with LLC tumor cells following microbial bioparticle treatment.** Two-photon microscopy was used to visualize Ly6G neutrophil (red) interacting with LLC cells (green) 24 hours after microbial bioparticle treatment. Magenta tracks indicate neutrophil paths. Projection of a time-lapse series imaged in eGFP-LLC tumors in the ear flap of BigRed/Catchup^IVM-red^ mice. Dimensions: 166 μm x 166 μm x 63 μm x 20 minutes. Images were taken 30 seconds apart. Elapsed time is shown as days:hours:min:seconds.

**Video 4. Neutrophil interactions with LLC cells lead to tumor cell blebbing.** Two-photon microscopy was used to visualize Ly6G neutrophil (red) interactions with LLC cells (green) 24 hours following microbial bioparticle treatment. Projection of a time-lapse series imaged in eGFP-LLC tumors in the ear flap of BigRed/Catchup^IVM-red^ mice. Example 1 dimensions: 125 μm x 125 μm x 93 μm x 60 minutes. Images were taken every 60 seconds. Example 2 dimensions: 125 μm x 125 μm x 78 μm x 60 minutes. Images were taken every 60 seconds. Elapsed time is shown as days:hours:min:seconds.
